## Supplementary material for "Conformational distributions of isolated myosin motor domains encode their mechanochemical properties": Supplimentary Information

### Supplement to *Structural dynamics of nucleotide-free myosin motor domains encode their duty ratios and ADP release rates*

Justin R. Porter, Artur Meller, Maxwell I. Zimmerman,
Michael J. Greenberg, and Gregory R. Bowman

### Contents


**1 Simulations details** **3**

**2 Markov State Model hyperparameters** **5**

**3 Further detail on P-loop PCA** **6**

**4 Structural and statistical characteristics of P-loop states** **9**

**5 Experimentally-measured biochemical properties** **10**

### 1 Simulations details

Each gene (listed in Table S1) was rendered to a 3D model by homology using SWISS-MODEL, then solvated and minimized, then equilibrated with position restraints on all heavy atoms, then simulated at fixed temperature and pressure. Most simulations were performed on folding@home [1], but MYH11 simulations were performed on our local cluster equipped with several different type of GPU-equipped computers (see below).

| Gene | Protein Name | Construct | Species | Template | Agg. Sim. [µs] |
|---|---|---|---|---|---|
| MYH13 | Extraocular | 4–781 | *H. sapiens* | 4PA0 [2] | 271.9 |
| MYH7 | $\beta$-cardiac | 2–780 | *H. sapiens* | 4PA0 [2] | 276.2 |
| MYH10 | Nonmuscle IIb-B2 | 8–791 | *H. sapiens* | 4PD3 [3] | 323.0 |
| MYO1B | Myosin-I | 5–703 | *H. sapiens* | 4L79 [4] | 282.3 |
| MYO5A | Myosin-Va | 2–762 | *H. sapiens* | 1W8J [5] | 297.5 |
| MYO6 | Myosin-VI | 2–770 | *H. sapiens* | 2BKI [6] | 295.0 |
| MYO7A | Myosin-VIIa | 3–742 | *H. sapiens* | 1OE9 [7] | 130.9 |
| MYO10 | Myosin-X | 3–740 | *H. sapiens* | 2AKA [8] | 126.2 |
| MYH11 | Chicken gizzard | wt/2–782 | *G. gallus* | 4PD3 [3] | 6.0 |
| MYH11 | Chicken gizzard | alanine | *G. gallus* | 4PD3 [3] | 6.4 |
| MYH11 | Chicken gizzard | xenopus | *G. gallus* | 4PD3 [3] | 16.5 |
| MYH11 | Chicken gizzard | $\Delta$loop 1 | *G. gallus* | 4PD3 [3] | 10.5 |

Table S1: *Summary of simulations performed for this study.* Gene names are those found in PubMed Gene for the appropriate organism, and residue numbers are those used in the given template.

Each system's energy was minimized using steepest descents until the maximum force on any atom decreased below 1000 $\mathrm{kJ\,mol^{-1}\,nm^{-1}}$ using a step size of 10 pm, a 1.2 nm cutoff for neighbor list, electrostatic interactions, and van der Waals interactions. Solvent was then relaxed at fixed pressure and temperature 300 K with the constraint of 1000 $\mathrm{kJ\,mol^{-1}\,nm^{-1}}$ applied to the protein heavy atoms and 2 fs as the time step. In this relaxation simulation, the temperature of the system is kept to 300 K using a Bussi-Parinello thermostat with a time constant of 0.1 ps [9]. The pressure was kept to 1 atm with an isotropic Parrinello-Rahman barostat with a relaxation time of 1 ps and a compressibility of $4.5 \times 10^{-5}\ \mathrm{bar^{-1}}$ [10]. A cut-off distance of 0.9 nm was used for the van der Waals and short-range electrostatic interactions. The Particle-Mesh-Ewald method was employed to recover the long-range electrostatic interactions with

0.12 nm as the grid spacing and with a fourth order spline [11]. All covalent bonds involving hydrogen were constrained using LINCS (Hess et al. 1997). After these equilibration runs, the restraints on heavy atoms were removed. Virtual sites were used to allow for a 4 fs time [12].

Simulations run on our local cluster included nodes with the following specifications:

1. Nodes with an Intel Xeon E5-2650 v2 CPU clocked at 2.60GHz, 16 GB RAM, and an nVidia Tesla K20 GPU.
2. Nodes with an Intel Xeon E5-2630 v3 CPU clocked at 2.40GHz, 16 GB RAM, and an nVidia Titan Xp GPU
3. Nodes with an Intel Xeon E5-2690 v4 CPU clocked at 2.60GHz, 32 GB RAM, and an nVidia Tesla P100 GPU
4. Nodes with an Intel Xeon Gold 6148 CPU clocked at 2.40GHz , 48 GB RAM, and an nVidia Quadro RTX 6000 GPU.

### 2 Markov State Model hyperparameters

| Simulation Set | No. of States | Cluster Radius [$nm^2$] | Lag Time [ns] |
|---|---:|---:|---:|
| MYH13 | 14102 | 7.4 | 0.4 |
| MYH7 | 5128 | 7.34 | 0.5 |
| MYH10 | 7746 | 8.0 | 1.5 |
| MYO1B | 6458 | 6.6 | 0.8 |
| MYO5A | 4728 | 7.25 | 0.4 |
| MYO6 | 4193 | 6.9 | 0.9 |
| MYO7A | 8737 | 6.9 | 0.4 |
| MYO10 | 9273 | 6.9 | 0.4 |
| MYH11, wild-type | 8050 | 4.9 | 1.5 |
| MYH11, alanine sub. | 7822 | 4.9 | 1.5 |
| MYH11, xenopus | 12804 | 5.2 | 1.5 |
| MYH11, $\Delta$loop 1 | 8925 | 5.0 | 1.5 |

Table S2: *Parameters of whole-motor Markov state models used in this study.* As explained in the *Methods* section of the main text, each trajectory set was converted to sidechain SASA, clustered using $k$-centers until the largest distance of any point to the nearest cluster center reached the stopping criterion ("Cluster Radius" above), and then refined with 5 rounds of $k$-medoids. Markov state models

### 3 Further detail on P-loop PCA

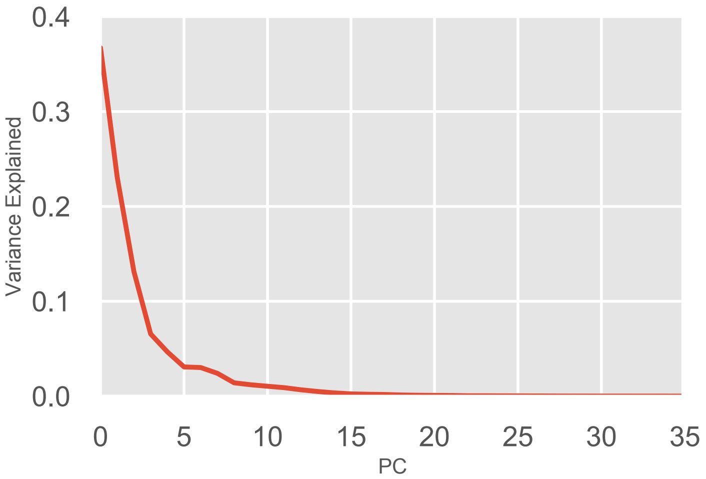


Figure S1: **Variance explained by each component of the PCA of the of the P-loop on MYH7.** Each Principal Component (PC) in the PCA captures a certain fraction of the variance of the input data. Each PC is ranked by the amount of variance it explains. The above figure plots the PC number against the fraction of the overall variance it explains. Because 36 distances were used in total, 36 principal components are able to capture 100% of the variance. The top four PCs were used in clustering and for $k$-nearest neighbor state assignment.

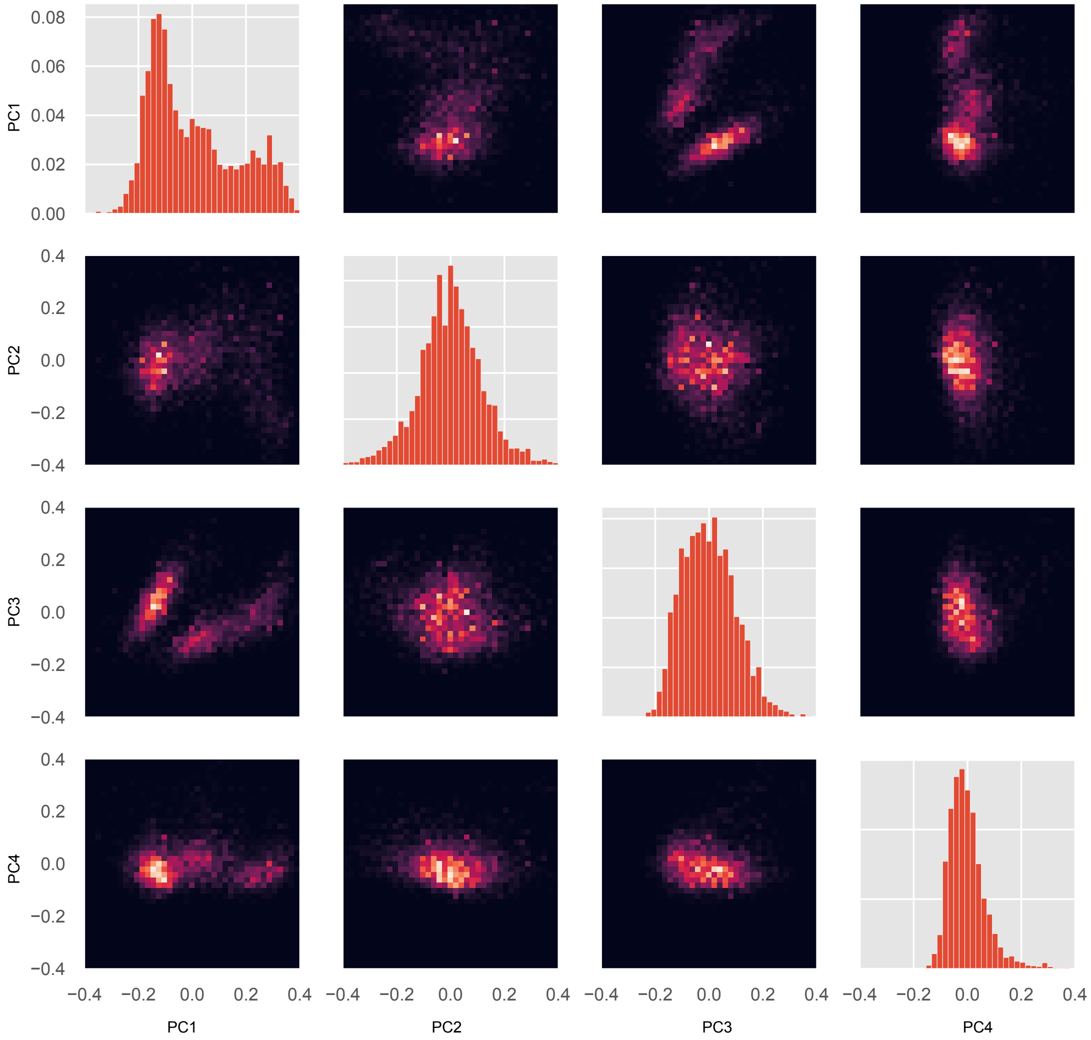


Figure S2: **Joint and marginal distributions for all pairs of PCs.** Each conformation in the whole-motor MSM has a value for each PC. Using these values and the probability for each conformation, a probability distribution for every individual PC's value, and for every pair of PCs values, were constructed. For MYH7, we visualized every marginal distribution (a single PC's probability distribution) and every pair of PCs' joint distribution. *Diagonal plots*, marginal distributions for PCs 1 (top left) through 4 (bottom right). *Off-diagonal plots*, joint distributions for all pairs of PCs; each column $i$ and row $j$ has PC $i$ on the $x$-axis and PC $j$ on the $y$-axis, respectively.

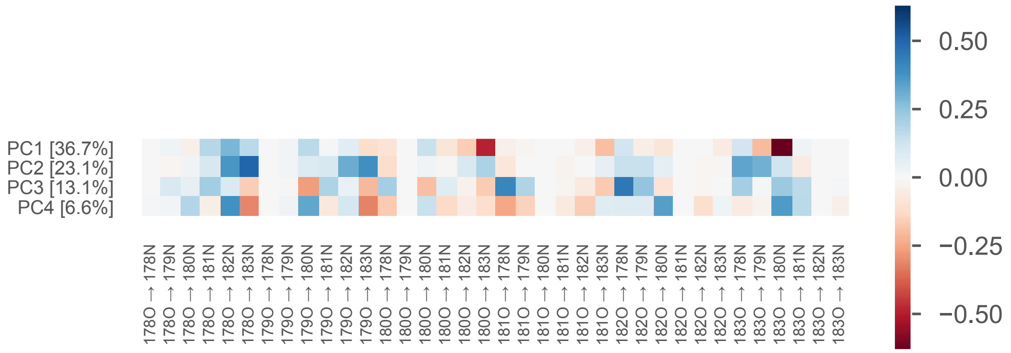


Figure S3: **Weights of the first four principle components of the P-loop on MYH7.** The value of each PC for a particular conformation is a linear combination of each of the 36 input distances. In other words, it is the dot product of the vector of distances and the vector representing the PC (which weights each distance differently). This figure shows the relative contribution of each distance ($x$-axis) to each PC ($y$-axis). Percentages following principle component (PC) index indicate the fraction of the variance explained by that PC. Labels on the $x$-axis indicate a distance from [residue number][atom name] to [residue number][atom name], using the residue numbering found in the MYH7 crystal model 4PA0 [2].

### 4 Structural and statistical characteristics of P-loop states

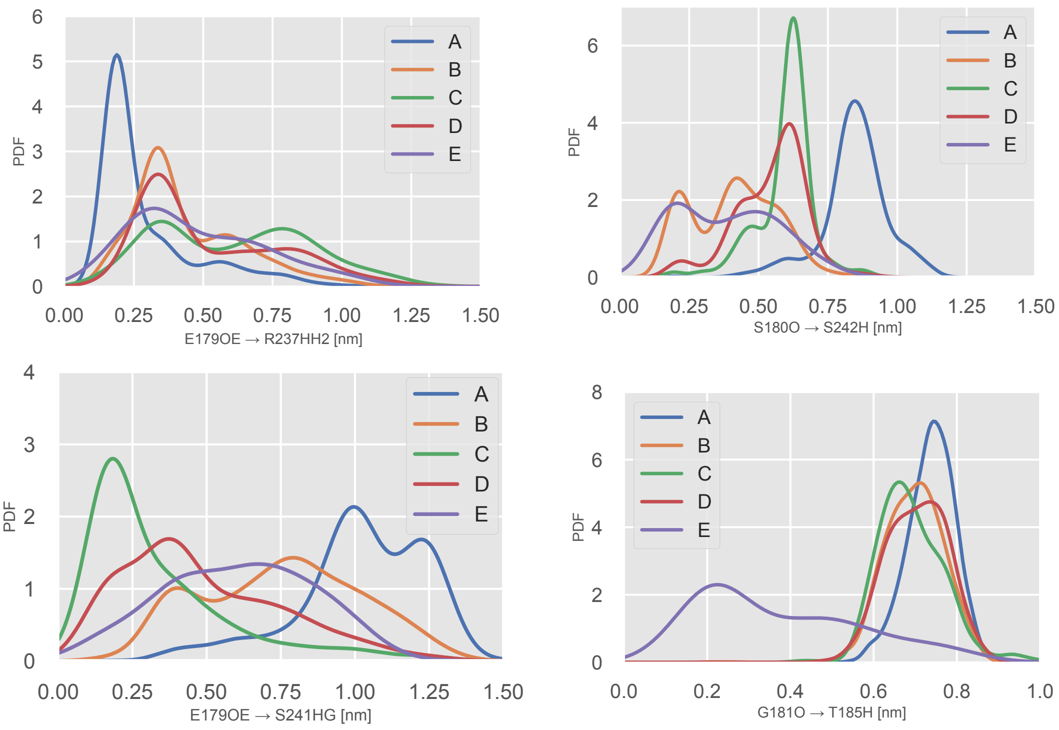


Figure S4: **Specific interactions with Switch-I residues are statistical hallmarks of P-loop states.** *Top left*, the PDF over the distance between E179 sidechain carbonyl oxygens to the sidechain of R237. An interaction is a hallmark of the A state. *Top right*, the PDF over the S180O to S242H backbone-backbone distance. Close values are a hallmark of states B and E. *Bottom left*, the PDF over the E179 sidechain carbonyl oxygen to S242 amide proton distance. Low values are a hallmark of state C. *Bottom left*, the PDF over the distance between the G181 backbone carbonyl oxygen and the T185 backbone amide proton. Low values are a hallmark of state E. This is the $i \rightarrow i+4$ interaction typical of an alpha-helix.

### 5 Experimentally-measured biochemical properties

| Protein | Duty Ratio | ADP Release Rate [$s^{-1}$] | Citation |
|---|---|---|---|
| MYH13 | 0.1 | 400 | Johnson et al. [13] |
| MYH7 | 0.1 | 59 | Johnson et al. [13] |
| MYH10 | 0.3 | 0.37 | Nagy et al. [14] |
| MYO1B | 0.05 | 2.1 | Lewis et al. [15] |
| MYO5A | 0.7 | 12 | De La Cruz et al. [16] |
| MYO6 | 0.9 | 5.6 | De La Cruz et al. [17] |
| MYO7A | 0.9 | 2.1 | Watanabe et al. [18] |
| MYO10 | 0.6 | 18 | Kovacs et al. [19] |
| MYH11, wild-type | 0.15 | 79 | Sweeney et al. [20] |
| MYH11, alanine sub. | 0.15 | 34 | Sweeney et al. [20] |
| MYH11, xenopus | 0.15 | 40 | Sweeney et al. [20] |
| MYH11, $\Delta$loop 1 | 0.15 | 13 | Sweeney et al. [20] |

Table S3: **Experimental values for kinetic and thermodynamic parameters of motor domains used in this study.**

### References


[1] M Shirts and V S Pande. COMPUTING: Screen Savers of the World Unite! *Science*, 290(5498):1903–1904, December 2000.

[2] Donald A Winkelmann, Eva Forgacs, Matthew T Miller, and Ann M Stock. Structural basis for drug-induced allosteric changes to human $\beta$-cardiac myosin motor activity. *Nat. Commun.*, 6(1):7974, August 2015.

[3] Stefan Münnich, Salma Pathan-Chhatbar, and Dietmar J Manstein. Crystal structure of the rigor-like human non-muscle myosin-2 motor domain. *FEBS Lett.*, 588(24):4754–4760, December 2014.

[4] Henry Shuman, Michael J Greenberg, Adam Zwolak, Tianming Lin, Charles V Sindelar, Roberto Dominguez, and E Michael Ostap. A vertebrate myosin-I structure reveals unique insights into myosin mechanochemical tuning. *Proc. Natl. Acad. Sci. U.S.A.*, 111(6):2116–2121, February 2014.

[5] Pierre-Damien Coureux, H Lee Sweeney, and Anne Houdusse. Three myosin V structures delineate essential features of chemo-mechanical transduction. *The EMBO Journal*, 23(23):4527–4537, November 2004.

[6] Julie Ménétrey, Amel Bahloul, Amber L Wells, Christopher M Yengo, Carl A Morris, H Lee Sweeney, and Anne Houdusse. The structure of the myosin VI motor reveals the mechanism of directionality reversal. *Nature*, 435(7043): 779–785, June 2005.

[7] Pierre-Damien Coureux, Amber L Wells, Julie Ménétrey, Christopher M Yengo, Carl A Morris, H Lee Sweeney, and Anne Houdusse. A structural state of the myosin V motor without bound nucleotide. *Nature*, 425(6956):419–423, September 2003.

[8] Thomas F Reubold, Susanne Eschenburg, Andreas Becker, Marilyn Leonard, Sandra L Schmid, Richard B Vallee, F Jon Kull, and Dietmar J Manstein. Crystal structure of the GTPase domain of rat dynamin 1. *Proc. Natl. Acad. Sci. U.S.A.*, 102(37):13093–13098, September 2005.

[9] Giovanni Bussi, Davide Donadio, and Michele Parrinello. Canonical sampling through velocity rescaling. *J. Chem. Phys.*, 126(1):014101, January 2007.

[10] Michele Parrinello and Aneesur Rahman. Polymorphic transitions in single

crystals: A new molecular dynamics method. *Journal of Applied physics*, 52 (12):7182–7190, 1981.

[11] Jiri Kolafa and John W Perram. Cutoff errors in the ewald summation formulae for point charge systems. *Molecular Simulation*, 9(5):351–368, 1992.

[12] K Anton Feenstra, Berk Hess, and Herman JC Berendsen. Improving efficiency of large time-scale molecular dynamics simulations of hydrogen-rich systems. *Journal of Computational Chemistry*, 20(8):786–798, 1999.

[13] Chloe A Johnson, Jonathan Walklate, Marina Svicevic, Srbolujub M Mijailovich, Carlos D Vera, Anastasia Karabina, Leslie A Leinwand, and Michael A Geeves. The ATPase cycle of human muscle myosin II Isoforms: adaptation of a single mechanochemical cycle for different physiological roles. *J. Biol. Chem.*, page jbc.RA119.009825, August 2019.

[14] Attila Nagy, Yasuharu Takagi, Neil Billington, Sara A Sun, Davin K T Hong, Earl Homsher, Aibing Wang, and James R Sellers. Kinetic characterization of nonmuscle myosin IIb at the single molecule level. *J. Biol. Chem.*, 288(1): 709–722, January 2013.

[15] John H Lewis, Michael J Greenberg, Joseph M Laakso, Henry Shuman, and E Michael Ostap. Calcium Regulation of Myosin-I Tension Sensing. *Biophys. J.*, 102(12):2799–2807, June 2012.

[16] Enrique M De La Cruz, Amber L Wells, Steven S Rosenfeld, E Michael Ostap, and H Lee Sweeney. The kinetic mechanism of myosin V. *Proc. Natl. Acad. Sci. U.S.A.*, 96(24):13726–13731, November 1999.

[17] Enrique M De La Cruz, E Michael Ostap, and H Lee Sweeney. Kinetic mechanism and regulation of myosin VI. *J. Biol. Chem.*, 276(34):32373–32381, August 2001.

[18] Shinya Watanabe, Reiko Ikebe, and Mitsuo Ikebe. Drosophila myosin VIIA is a high duty ratio motor with a unique kinetic mechanism. *J. Biol. Chem.*, 281 (11):7151–7160, March 2006.

[19] Mihaly Kovacs, Fei Wang, and James R Sellers. Mechanism of action of myosin X, a membrane-associated molecular motor. *J. Biol. Chem.*, 280(15): 15071–15083, April 2005.

[20] H Lee Sweeney, Steven S Rosenfeld, Fred Brown, Lynn Faust, Joe Smith, Jun Xing, Leonard A Stein, and James R Sellers. Kinetic Tuning of Myosin via a

Flexible Loop Adjacent to the Nucleotide Binding Pocket. *J. Biol. Chem.*, 273 (11):6262–6270, March 1998.
